# Mapping the Fascicular Morphology and Organization of the Human Sciatic Nerve via High-Resolution MicroCT Imaging

**DOI:** 10.1101/2025.06.28.662133

**Authors:** Jichu Zhang, Valerie H. Lam, Noa B. Nuzov, Brandon A. S. Brunsman, Tatiana Pascol, Ashley Onabiyi, Rebecca Prince, Havisha Kalpatthi, Kenneth Gustafson, Ronald Triolo, Nicole Pelot, Andrew Crofton, Andrew Shoffstall

## Abstract

**Objective:** Implanted neuroprostheses can restore standing and walking after spinal cord injury and somatosensation after limb loss. Yet current approaches often fail to reliably activate hamstring muscles crucial for upright stability and mobility or to target afferent fibers for sensory restoration. We developed a novel methodology using high-resolution micro-computed tomography (microCT) to visualize and track fascicle groups innervating distinct hamstring muscles along the human sciatic nerve. This approach provides a framework for mapping fascicular topography in complex neural pathways to optimize standing neuroprostheses.

**Methods:** Bilateral sciatic nerves were dissected and excised from an embalmed human cadaver, annotated with branch names, and stained with phosphotungstic acid before undergoing microCT scanning at 11.4 μm isotropic resolution. Images were segmented with a 3D U-Net convolutional neural network. Segmentation results were used to quantify morphological metrics and track fascicular organization along ∼25 cm of the nerve. MicroCT reconstructions were validated against histological cross sections.

**Results:** Gross dissection revealed matched proximal-to-distal branching between left and right sciatic nerves: branch to long head of the biceps femoris (lhBF), branch to hamstring part of the adductor magnus and semimembranosus (HAM/SM), and branch to semitendinosus (ST). All branches originated medially and followed an inferomedial trajectory. Branch-free lengths of the sciatic exhibited asymmetry, especially between the lumbosacral roots to the first branch (5.5 cm left vs. 1.5 cm right) and lhBF to the HAM/SM branch (9.0 cm left vs. 16.5 cm right). MicroCT analysis revealed bilateral symmetry in fascicle diameters (∼0.4 mm) and total fascicle counts (∼84) but asymmetry in hamstring-innervating fascicle counts (left ∼7, right: ∼9). The 3D fascicular maps revealed that hamstring fascicles were located in the anteromedial portion of the sciatic nerve cross section and remained separate for distances up to 15.9 cm proximal to their branching points.

**Significance:** Our microCT-based approach enables efficient, high-resolution 3D mapping of fascicular organization within large, complex peripheral nerves like the sciatic nerve, overcoming previous technical limitations. This methodology informs development of neuroprostheses with improved hamstring muscle activation for enhanced standing function.

## 1. Introduction

Implanted motor system neuroprostheses offer significant functional improvements for individuals with spinal cord injuries (SCI) by electrically stimulating peripheral nerves to activate the otherwise paralyzed muscles (Kilgore et al., 2023; Peckham and Knutson, 2005). Neuroprostheses aim to deliver selective, coordinated stimulating currents to generate useful movements in targeted muscle groups (Peckham and Knutson, 2005). Studies have shown that neuroprostheses can support basic activities, such as standing and walking, following paraplegia (Uhlir et al., 2000; Hardin et al., 2007; Fisher et al., 2008; Triolo et al., 2012; Davis et al., 2001). Such functional restoration addresses a critical need for people with SCI, who rank the recovery of lower extremity mobility among their highest rehabilitative goals (Brown-Triolo et al., 2002; Anderson, 2004). Beyond functional gains, these systems improve independence in conducting activities of daily living (Kilgore et al., 2023) and enhance quality of life (Rohde et al., 2012). The upright posture enabled by standing neuroprostheses mitigates complications and risk factors caused by immobility after SCI—including osteoporosis, pressure ulcers, and muscle spasticity (Sezer et al., 2015)—and facilitates transfers between surfaces (Davis et al., 2001; Triolo et al., 2012).

Despite these benefits, current neuroprostheses cannot consistently provide prolonged, independent standing (Fisher et al., 2008). Individuals experience short standing duration (Mushahwar et al., 2007), muscle fatigue (Kobetic et al., 1997), inadequate body weight support, and variations in clinical outcomes between recipients (Fisher et al., 2008). These limitations partly stem from incomplete and inconsistent activation of muscle groups responsible for stabilizing the hips. Existing systems (Davis et al., 2001; Fisher et al., 2008; Triolo et al., 2012; Uhlir et al., 2000) rely on knee extension (via quadriceps), trunk stabilization (via erector spinae), and modest hip extension (via the gluteus maximus) but they do not provide sufficient activation of the hamstring muscle group (Fisher et al., 2008; Schiefer et al., 2008), which comprises the semitendinosus (ST), semimembranosus (SM), and biceps femoris (BF), along with the functionally similar hamstring part of adductor magnus (HAM). These muscles—hereafter termed “hamstrings”—provide strong hip extension and stabilize the pelvis during standing (Kuo and Zajac, 1993), which are necessary for neuroprostheses to maintain an erect posture and reduce dependence on arm support (Kobetic et al., 1999). Improved activation of hamstrings would therefore address key limitations of current standing systems by enhancing postural stability and extending standing duration through more biomechanically efficient weight support (Kuo and Zajac, 1993). In stepping and transfer applications, enhanced hamstring activation may provide additional benefits by decelerating knee extension to prevent hyperextension at terminal swing, stabilizing pelvic posture, advancing the body during stance, and—via the short head of the biceps femoris—facilitating knee flexion for foot clearance in early swing.

The hamstrings are innervated by the sciatic nerve, the longest and largest nerve in the human body (Giuffre et al., 2023). The sciatic originates from the L4–S3 spinal nerves; it traverses the posterior thigh, spans the length of the leg, and bifurcates into two branches that extends toward the foot (Giuffre et al., 2023). The nerve is ∼30 cm in length in humans from the greater trochanter to its bifurcation into the tibial and common fibular nerves in the popliteal fossa (Gustafson et al., 2012). For clarity, hereafter, we use the term “sciatic” to refer to the nerve from its formation at the lumbosacral roots to its bifurcation.

Given the extensive innervation provided by the sciatic nerve, controlling the selective activation of targeted fibers is important for effective lower limb motor neuroprostheses (Grill et al., 2009), as well as sensory neuroprostheses after lower limb loss (Charkhkar et al, 2018). Anatomically realistic computational models can be leveraged to design multi-contact electrodes and stimulation parameters that achieve selective activation of targeted fibers. For example, computational models guided the development of neuroprostheses that selectively control knee extensors and hip flexors via the femoral nerve (Schiefer et al., 2010, 2008), as well as plantar and dorsiflexors via the tibial and common fibular nerves, respectively (Schiefer et al., 2013, 2012).

Model-based design of selective neuroprostheses requires appropriate anatomical inputs, including the nerve diameter, fascicular structure, and functional organization, which is particularly challenging given the size and complexity of the human sciatic nerve, which contains up to 70 fascicles (Sladjana et al., 2008). Functional mapping studies of peripheral nerves typically use histology with serial sectioning or manual tracing of fascicles through microdissection. However, these approaches are only tractable for smaller nerves across shorter distances (e.g., 2–5 cm, ≤ 20 fascicles) (Gustafson et al., 2012, 2009), and such brute-force methods fail to scale to the sciatic nerve. The nerve’s extended length demands an extremely large number of sections, and its fascicular complexity makes tracking across sparse 2D cross sections both labor-intensive and prone to error (Gustafson et al., 2012). Indeed, prior studies of the sciatic nerve examined its general somatotopic organization for restricted sample counts and anatomical spans (Gustafson et al., 2012; McKinley, 1921; Sunderland and Ray, 1948), but yielded incomplete fascicular mapping, particularly for the proximal segments critical to hamstring control.

Alternative imaging modalities for fascicular mapping are also limited in spatial resolution and/or field of view. Magnetic resonance neurography provides ∼0.5 mm spatial resolution, which is insufficient to visualize fine fascicular structures (Soldatos et al., 2013); high-resolution ultrasound at 22 MHz resolves only ∼50% of sciatic fascicles due to its ∼0.2 mm lateral resolution (Snoj et al., 2024); magnetic resonance microscopy improves resolution to ∼50 μm, but it is restricted to ∼2.5 cm nerve segments (Yao et al., 2018). These technical constraints highlight the need for an improved method for mapping the sciatic nerve. A new approach with micron-level resolution spanning the nerve’s full length is essential to provide the anatomical data needed for designing lower limb neuroprostheses.

Micro-computed tomography (microCT) offers a novel solution to this methodological barrier by providing three-dimensional (3D) X-ray imaging of biological samples (Stauber and Müller, 2008) with both high spatial resolution (3.3–50 μm) and a multi-centimeter field of view. Though widely used for skeletal imaging, microCT can also effectively visualize soft tissues, including neural structures, when paired with contrast agents (Descamps et al., 2014; Stauber and Müller, 2008). Recent studies have used microCT to image fascicular anatomy in rat sciatic nerve, pig vagus nerve, and human vagus nerve after staining with Lugol’s iodine, osmium tetroxide, or phosphotungstic acid (PTA) (Sladjana et al., 2008; Upadhye et al., 2022; Thompson et al., 2020; Buyukcelik et al., 2023; Thompson et al., 2023; Rossetti et al., 2025; Upadhye et al., 2025a; Zhang et al., 2026). Osmium tetroxide staining has also enabled quantitative analysis of pig spinal rootlets based on microCT images (Chin et al., 2024).

Here, we demonstrate the first application of microCT for mapping the morphology and functional organization of fascicles in the human sciatic nerve. We focused on fascicles innervating muscles that are critical for standing function (ST, SM, long head of BF [lhBF], and HAM). Through detailed cadaveric dissection and PTA contrast enhancement, we achieved visualization of fascicular structure at 11.4 μm isotropic resolution. We automated the segmentation of the fascicular structure using a 3D convolutional neural network (Zhang et al., 2026) and analyzed the resulting morphology and functional organization. This approach enables continuous mapping of fascicles innervating the hamstring muscles from their distal branching points to the lumbosacral roots, overcoming previous limitations in resolution and field of view. Our methods for generating the neuroanatomical map of the human sciatic nerve provide the data required for computational optimization of neural interface designs for robust, selective, and consistent hamstring activation and improved lower limb rehabilitation via neuroprostheses.

## 2. Methods

We adopted the dissection, tissue handling, and imaging pipeline developed for cadaveric human vagus nerves (Pelot et al., 2025).

### 2.1. Cadaver Preparation

The left and right sciatic nerves with their branches were harvested from a 41-year-old Black male. The cadaver was obtained through the Anatomical Gift Program at Case Western Reserve University (CWRU), Cleveland, OH. This study was determined to be exempt by CWRU’s Institutional Review Board because it involved de-identified cadaveric tissue, and no protected health information was collected. The cadaver was embalmed with a traditional formaldehyde-phenol embalming solution containing isopropanol (0.77%), methanol (1.74%), formaldehyde (3.9%), phenol (4.88%), propylene glycol (5.05%), ethanol (13.85%), and water (69.81%). The embalming fluid was administered via injection into the right femoral artery with supplemental embalming fluid injected subcutaneously 1 week later in areas with inadequate embalming, as indicated by tissue suppleness and/or lack of discoloration, including in the torso, gluteal region, and forearms.

### 2.2. Anatomical Dissection

#### 2.2.1. Dissection

The dissection was performed with the body prone. Skin, connective tissue, and gluteal muscles were removed to isolate the lumbosacral roots and the sciatic nerves as they exited the greater sciatic foramina. The dissection continued inferiorly, where the surrounding fascia and vasculature were removed to isolate nerve branches and follow them to their muscle targets. The lhBF was detached distally near the knee and reflected for improved visibility of the nerve; all other hamstring muscles (SM, ST, short head of BF [shBF], HAM) remained attached both proximally and distally. On both left and right sides, sciatic branches innervating the hamstrings and their muscle targets were identified and documented (Figure 1). Throughout the dissection, the nerve was periodically sprayed with a wetting solution (40% commercial bleach, 59.99% water, and 0.01% phenol) to maintain tissue hydration, prevent desiccation and inhibit microbial growth.

**Figure 1.**
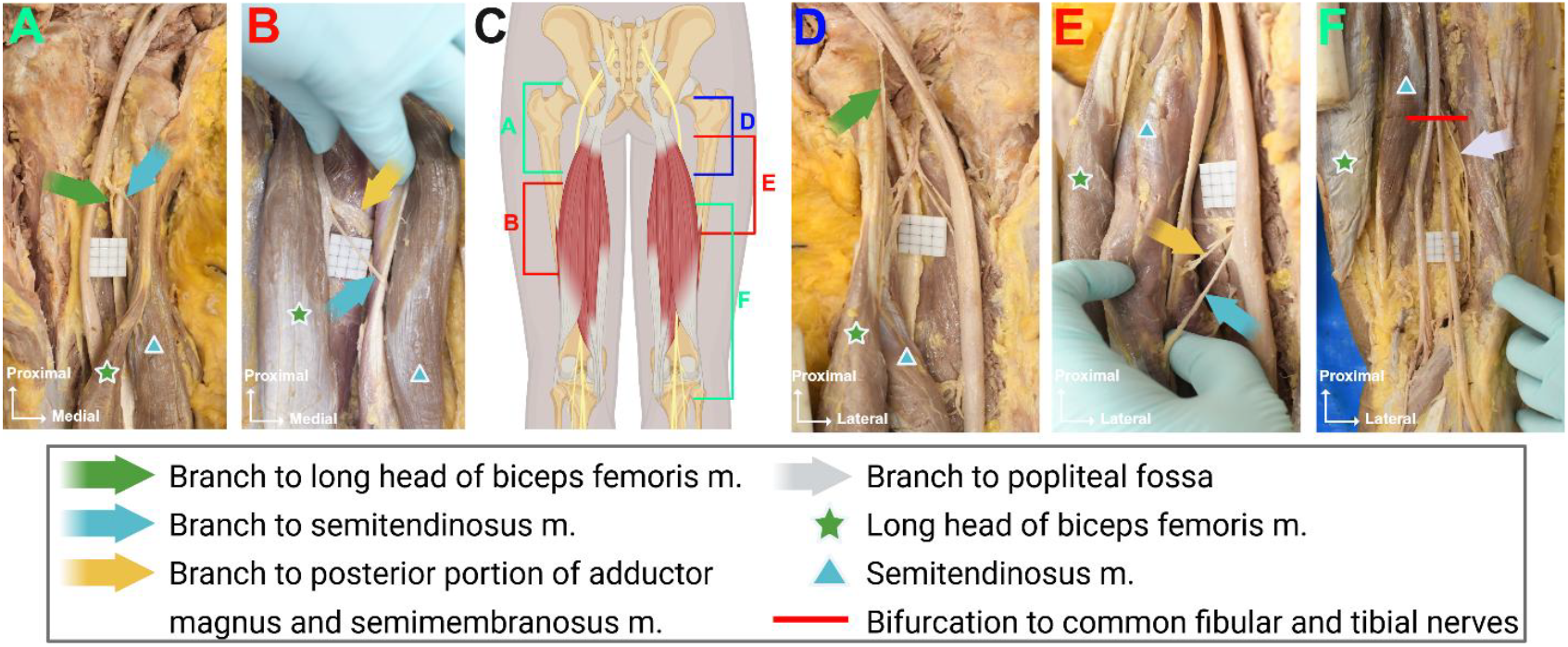
*In situ* photos of bilaterally dissected sciatic nerves, branches that innervate the hamstrings, and hamstring muscles. Left (A-B) and right (D-F) sciatic nerves are viewed posteriorly with the body prone. The right lhBF tendon was cut distally and its muscle belly reflected medially to facilitate visualization of the right sciatic nerve. (C) Schematic diagram showing the relative positions of the photos. (A) Left sciatic nerve emerging from the pelvis with branches to ST and lhBF muscles, and (B) branches to HAM, SM, and ST muscles. (D) Right sciatic nerve emerging from the pelvis with a branch innervating the lhBF (cut), and (E) branches to HAM, SM, and ST. (F) Bifurcation of the sciatic nerve into the common fibular and tibial nerves, and the branch traveling to the PF. Scale: 0.5 cm x 0.5 cm squares etched into a 2 cm x 2 cm board. m., muscle; lhBF, long head of biceps femoris; HAM, hamstring part of the adductor magnus; PF, popliteal fossa; shBF, short head of biceps femoris; SM, semimembranosus; ST, semitendinosus.

#### 2.2.2. Anatomical Annotations and Gross Measurements

The branch patterns of the right and left sciatic nerves were documented, including locations of branch points and target muscles. Branches were categorized based on their site of origin on the nerve (anterior, posterior, medial, or lateral) and their path relative to the sciatic. Branches were painted to assist with visual identification. To provide anatomical context, key skeletal landmarks were identified along the sciatic nerve, and their corresponding axial levels were marked directly on the sciatic with paint. These levels were: formation of the sciatic from lumbosacral roots, greater trochanter of the femur, lesser trochanter of the femur, and points at 7, 14, and 21 cm inferior to the lesser trochanter.

Gross anatomic measurements of the sciatic were performed bilaterally. The medial-lateral and anterior-posterior diameters of the sciatic were measured using digital calipers (VWR, Radnor, PA; precision of 0.1 mm) at the six marked anatomical levels. Following dissection and measurements, the sciatic was excised by transecting it proximally at the greater sciatic foramen and distally at its terminal branches. Then, the nerve was glued (Gorilla Super Glue Brush and Nozzle, Cincinnati, OH) to 3 cm x 9 cm (width x height) acrylic boards etched with 0.5 cm x 0.5 cm reference squares with the anterior surface of the nerve facing the board. The orientation of the origin of each branch was preserved when gluing the nerves to the boards, but branch trajectories were not reliably maintained due to limited space on the boards. Glue was only placed at the cut ends of the nerve and its branches. Branch-free length was defined as the linear distance longitudinally along the sciatic between consecutive branches as measured from the inferior margin of a proximal branch point to the superior margin of a distal branch point. Branch locations were approximated to the nearest 5 mm grid mark according to the excised nerve in post-removal photos.

### 2.3. MicroCT Imaging

#### 2.3.1. Tissue Staining

We stained the nerve with PTA to provide contrast between the fascicles and surrounding epineurium (Upadhye et al., 2025a, 2025c). A 3% phosphotungstic acid (PTA) solution was prepared by diluting a 10% stock PTA solution (HT152-250ML, Sigma-Aldrich, St. Louis, MO) with deionized water. The nerve was submerged in 1.5 L of this PTA staining solution and placed on an orbital shaker (SI-M1500, Scientific Industries, Bohemia, NY) set at 50 RPM for 96 hours. The prolonged staining duration ensured adequate penetration into the thick sciatic nerves. Staining quality was verified by examining tissue contrast in preliminary scans. Prior to microCT scanning, the nerve was covered with gauze saturated with 1x phosphate buffered solution (PBS, BP399-1, Fisher Scientific, Hampton, NH) and stored in an airtight container at 4°C to prevent dehydration.

#### 2.3.2. Scanning and Image Processing

We adopted a microCT protocol used to image human vagus nerves (Upadhye et al., 2025b). The stained nerve was cut at the end of each 9 cm-long acrylic board to create smaller segments that fit in the sample holder and the scanner’s sample tube. Each nerve segment on an acrylic board was placed in a cylindrical sample holder (35.4 mm diameter, 110 mm long). All microCT scans were performed using a μCT100 cabinet scanner (SCANCO Medical AG, Wangen-Brüttisellen, Switzerland) with the following parameters: 55 kV X-ray voltage, 145 μA current, 500 ms integration time, and 0.5 mm aluminum filter. The field of view was set at 35.2 mm, producing images with an isotropic resolution of 11.4 μm per pixel. Each scan was exported as a series of 16-bit DICOM slices (3072 x 3072). For efficient file storage and analysis, a volumetric image in OME-Zarr format (Moore et al., 2023) was created for each nerve sample at the original resolution.

### 2.4. Image Segmentation

We fine-tuned a 3D U-Net model, originally developed and pre-trained on 100 microCT images of PTA-stained human vagus nerves (Zhang et al., 2026) to segment similar images of human sciatic nerves.

#### 2.4.1. Ground Truth Development

Fascicle and epineurium boundaries in each microCT sub-volume (64 x 1536 x 3072) were manually outlined using 3D Slicer v5.6.1 (Fedorov et al., 2012) to serve as ground truth. The locations of nine manual segmentations (each 64 slices) were selected from regions where the pre-trained vagus nerve model (Zhang et al., 2026) showed suboptimal performance. The “Fill between slices” function in the “Segmentation” module was used to interpolate between manually segmented slices for fascicles with consistent locations and shapes along the nerve. Regions of fascicle splitting or merging were segmented manually. The interpolated manual segmentations were validated against the raw microCT image to ensure accurate boundaries and handling of fascicle branching. The resulting fascicle and epineurium masks were combined into a single 8-bit multiclass label matching the original voxel spacing.

#### 2.4.2. Neural Network Architecture

A 3D convolutional neural network based on the widely used U-Net architecture (Çiçek et al., 2016; Zhang et al., 2026) was used for automatic segmentation of fascicles and epineurium from microCT images of the sciatic nerve. The model network processed single-channel grayscale 3D volumes and output segmentation maps for three classes: background, fascicles, and epineurium. The network featured a 7-stage encoder-decoder structure. The number of features per stage was 32, 64, 128, 256, 320, 320, and 320. Each stage included two 3 x 3 x 3 convolutions, and residual connections were incorporated within each encoder stage. The decoder path mirrored the encoder, with skip connections from corresponding stages. Instance normalization and leaky rectified linear unit (ReLU) were used throughout the network. No dropout was used.

#### 2.4.3. Network Training

The network underwent a two-stage training process: initial pre-training on 100 PTA-stained microCT images (64 x 1536 x 3072) of human cervical and thoracic vagus nerve, acquired using identical scanner parameters (Zhang et al., 2026), followed by fine-tuning on an additional set of nine sciatic nerve sub-volumes collected for this study (Section 2.4.1). The nine sciatic nerve sub-volumes used for fine-tuning were selected from regions of the sciatic nerve images where the initial pre-trained model produced suboptimal prediction results. This transfer learning approach leveraged features learned from similar peripheral nerve structures and enabled efficient model adaptation with limited sciatic nerve training data.

The same training protocol was used in both pre-training and fine-tuning. The model was trained on randomly sampled 32 x 256 x 256 patches from the original volumes for 500 epochs with a batch size of 4. The network was optimized using stochastic gradient descent with Nesterov momentum (*μ* = 0.99), an initial learning rate of 0.01, and polynomial learning rate decay. To prevent overfitting given a small training dataset, a comprehensive set of data augmentation techniques was applied during training. Detailed specifications of the augmentation strategies are provided in Supplementary Note 1. All training was performed on an NVIDIA RTX A6000 GPU (Santa Clara, CA) with 48 GB of memory using 16-bit automatic mixed precision.

#### 2.4.4. Segmentation Performance and Prediction

The model showed high segmentation performance with a mean intersection over union (IoU) of 0.88 for fascicles and 0.92 for epineurium on in-training validation image patches. Following successful validation, the trained network was applied to generate segmentations of fascicles and epineurium for all acquired microCT scans. Each automated segmentation underwent visual inspection to verify anatomical accuracy. The quality-controlled segmentations were used for subsequent quantitative analyses.

### 2.5. Quantitative Analysis

#### 2.5.1. Morphological Measurements

Nerve and fascicle morphological metrics were measured at the original resolution from the segmented microCT images. Measurements were performed on cross sections along the nerve length longitudinally with the original slice spacing (11.4 μm): the number of fascicles, the effective diameters of individual fascicles, and the effective diameter of the nerve. The effective diameter was defined as 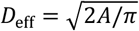, where *A* is the cross-sectional area of an individual fascicle or whole nerve. In addition, Feret diameters of the nerve cross section were measured in the vertical and horizontal directions to validate gross measurements of the anterior-posterior and medial-lateral diameters, respectively. All morphological measurements were calculated using custom Python (version 3.10) scripts and connected components algorithms from the OpenCV package (Bradski, 2000).

Branches identified during gross dissection were verified using microCT imaging. Vascular branches that lacked fascicles were excluded from analyses.

#### 2.5.2. Fascicle Tracking

Following segmentation, fascicles were tracked longitudinally along the nerve by matching fascicles in adjacent cross sections. The algorithm first identified individual fascicle instances from two segmented image frames as connected components. Then, spatial overlap between each potential fascicle pair across frames was calculated using a confusion matrix. One-to-one connections were established when a fascicle in one frame had significant overlap (IoU > 0.85) with exactly one fascicle in the adjacent frame. The threshold was determined empirically through validation with manual tracking.

The algorithm also handled splitting and merging of fascicles. A splitting event was defined as a single fascicle in the proximal frame dividing into two or more fascicles distally, while a merging event occurred when two or more proximal fascicles converged into a single fascicle distally. These events were identified when one fascicle (the parent) had an overlap with multiple child fascicles. To confirm these events, the algorithm verified area conservation by comparing the total cross-sectional area of the child fascicles with the parent fascicle area, allowing for a tolerance of ±15%, as fascicle area should be approximately conserved during splitting or merging (Upadhye et al., 2022). The algorithm was implemented in Python using OpenCV (Bradski, 2000) and scikit-image (Walt et al., 2014).

Fascicles of each hamstring-innervating branch were identified in the microCT segmentation, where the epineurium of the branch and the sciatic were distinct. Based on the tracking data, these hamstring fascicles were automatically backtracked from distal to proximal regions of the nerve. If fascicles innervating different target muscles shared a parent fascicle at any point during backtracking, the parent fascicle was classified as a “mixed hamstring fascicle”. The distance between each hamstring muscle’s branch point and the location where its fascicle group mixed with other fascicles for the first time was measured and recorded for each target muscle.

## 3. Results

### 3.1. Gross Anatomy of Sciatic Nerve and Hamstring Branches

The left and right sciatic nerves innervated the hamstring muscles in a mostly consistent proximal-to-distal order: lhBF, HAM/SM, and ST (Figure 2). Most hamstring muscles received one branch, except for the left ST, which received an additional sub-branch of the lhBF branch. Also, the SM branch came off the HAM branch on both sides. All hamstring branches emerged from the medial aspect of the sciatic nerve proximal to its bifurcation and followed an inferomedial trajectory to their target muscles.

**Figure 2.**
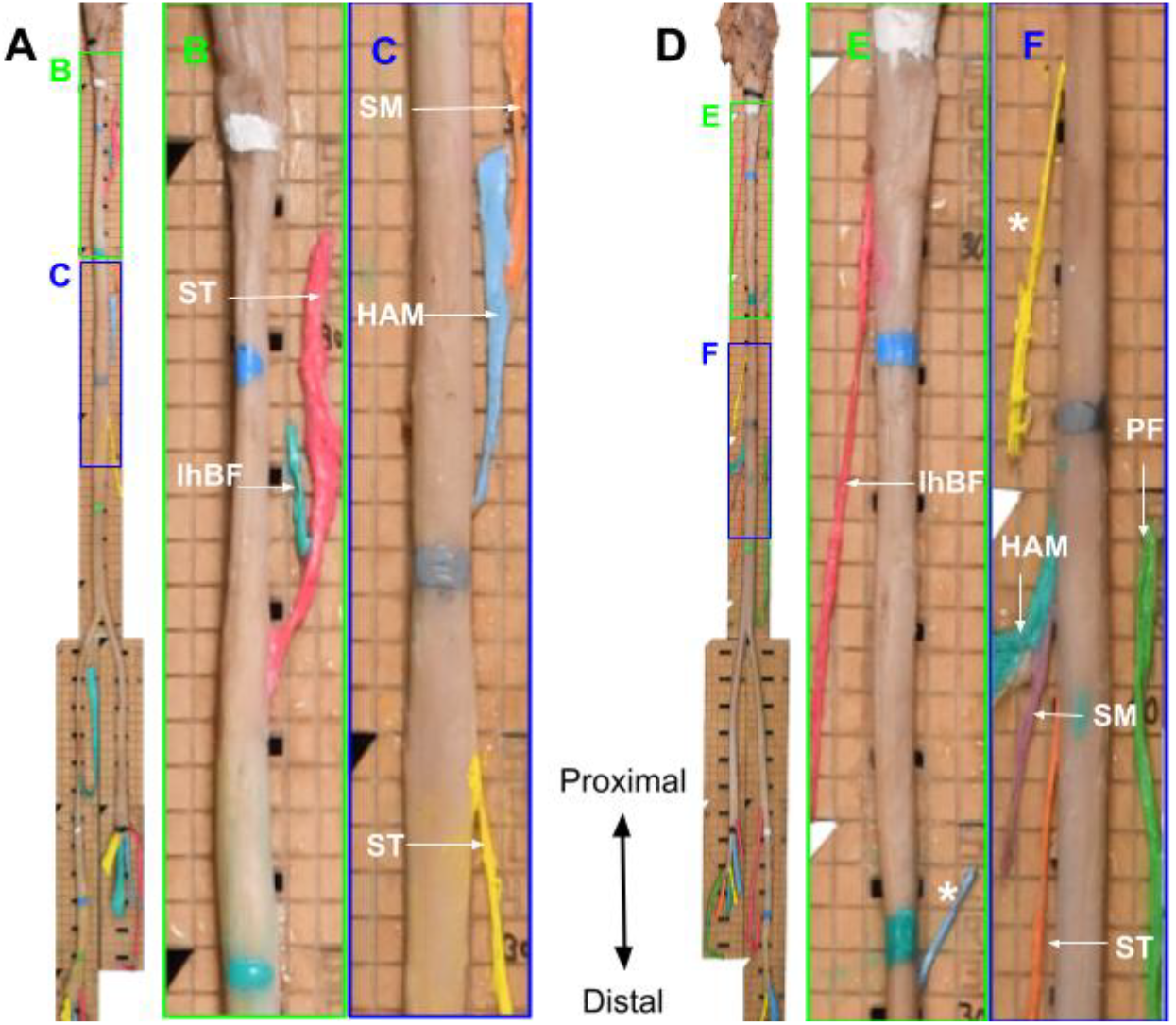
Excised left (A-C) and right (D-F) human sciatic nerves with labeled hamstring branches. Photos of the left (A) and right (D) sciatic nerves have color-coded boxes highlighting magnified regions with hamstring-innervating branches for the left (B, C) and right (E, F) sides. Labeled white arrows indicate muscle targets of dissected branches. Horizontal lines of paint on the sciatic indicate anatomical landmarks: white paint marks the sciatic formation from lumbosacral roots, blue paint marks the level of the lesser trochanter of the femur, teal paint marks 7 cm inferior to the lesser trochanter, and gray paint marks 14 cm inferior to the lesser trochanter. The etched grids on the backing boards are 0.5 cm x 0.5 cm. *, blood vessels; lhBF, long head of biceps femoris; HAM, hamstring part of the adductor magnus; PF, popliteal fossa; shBF, short head of biceps femoris; SM, semimembranosus; ST, semitendinosus.

Despite a relatively consistent pattern of hamstring nerve branching along the sciatic, the branch-free lengths between branch points were different between the left and right sides (Table 1). The most proximal segment, from lumbosacral roots to the first branch (lhBF), was 3.7 times longer on the left side (5.5 cm) than on the right (1.5 cm). The longest branch-free segment on both sides occurred between lhBF and HAM/SM branches, measuring 9.0 cm on the left and 16.5 cm on the right.

**Table 1.**
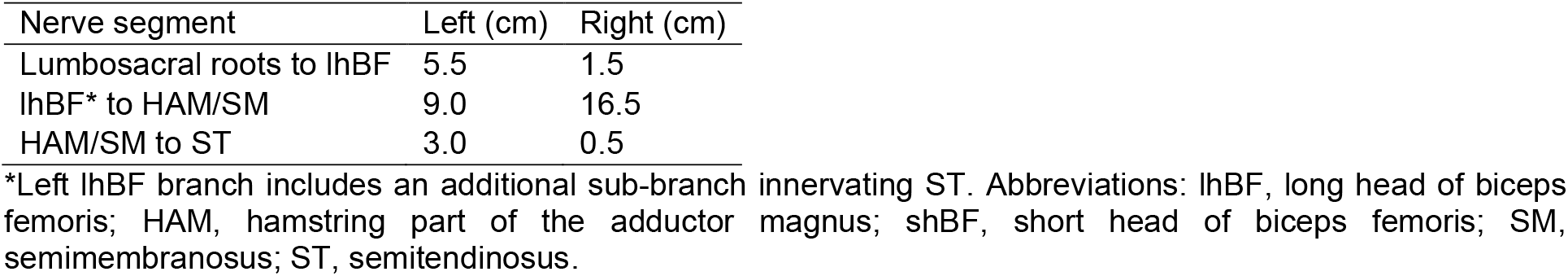
Branch-free lengths of bilateral sciatic nerves.

The sciatic nerve maintained an elliptical shape throughout its course, with a mean aspect ratio (medial-lateral/anterior-posterior) of 1.5 on the left side and 1.8 on the right (Figure 3). At the level of the lesser trochanter of the femur, the sciatic approached a more circular shape with aspect ratios of 0.9 (left) and 1.2 (right). Maximum medial-lateral diameters (11.7 mm) occurred at the level of the greater trochanter of the femur on both sides (Table S1).

**Figure 3.**
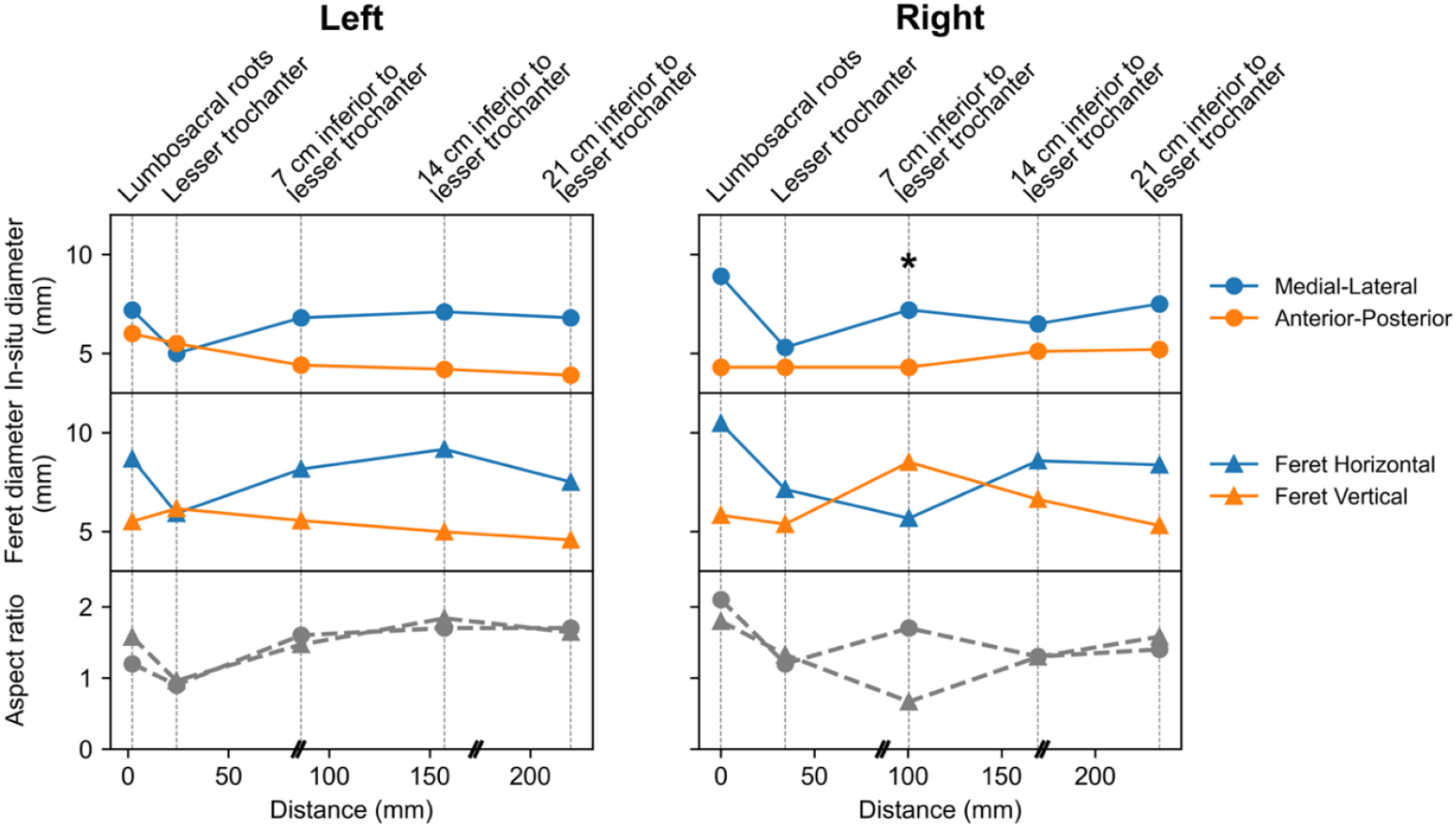
In situ and microCT-derived nerve diameter measurements. Feret diameters measured in the horizontal and vertical directions correspond to the medial-lateral and anterior-posterior nerve diameters measured in situ, respectively. The asterisk (*) indicates a location where rotation of the nerve on the backing board during staining and/or handling caused the horizontal and vertical Feret diameter directions to swap relative to the in situ directions. Measurements at the greater trochanter are reported in Table S1. Vertical dotted lines indicate the levels of anatomical landmarks. Double-slash marks (//) on the x-axis indicate transitions between microCT scanning grids.

### 3.2. MicroCT Characterization of the Fascicular Morphology of the Sciatic Nerve

MicroCT imaging enabled effective visualization of the detailed 3D fascicular structure of PTA-stained sciatic nerves and its hamstring branches across ∼30 cm length on each side (Figure 4). Fascicles with diameters from 0.1 to 0.7 mm were accurately segmented within the sciatic nerves and their branches. The documented branch patterns enabled precise identification of specific hamstring branches and their fascicles in the microCT images. The cross-sectional morphology in microCT images was verified against histological slides with H&E staining from the corresponding location (Supplementary Note 3).

**Figure 4.**
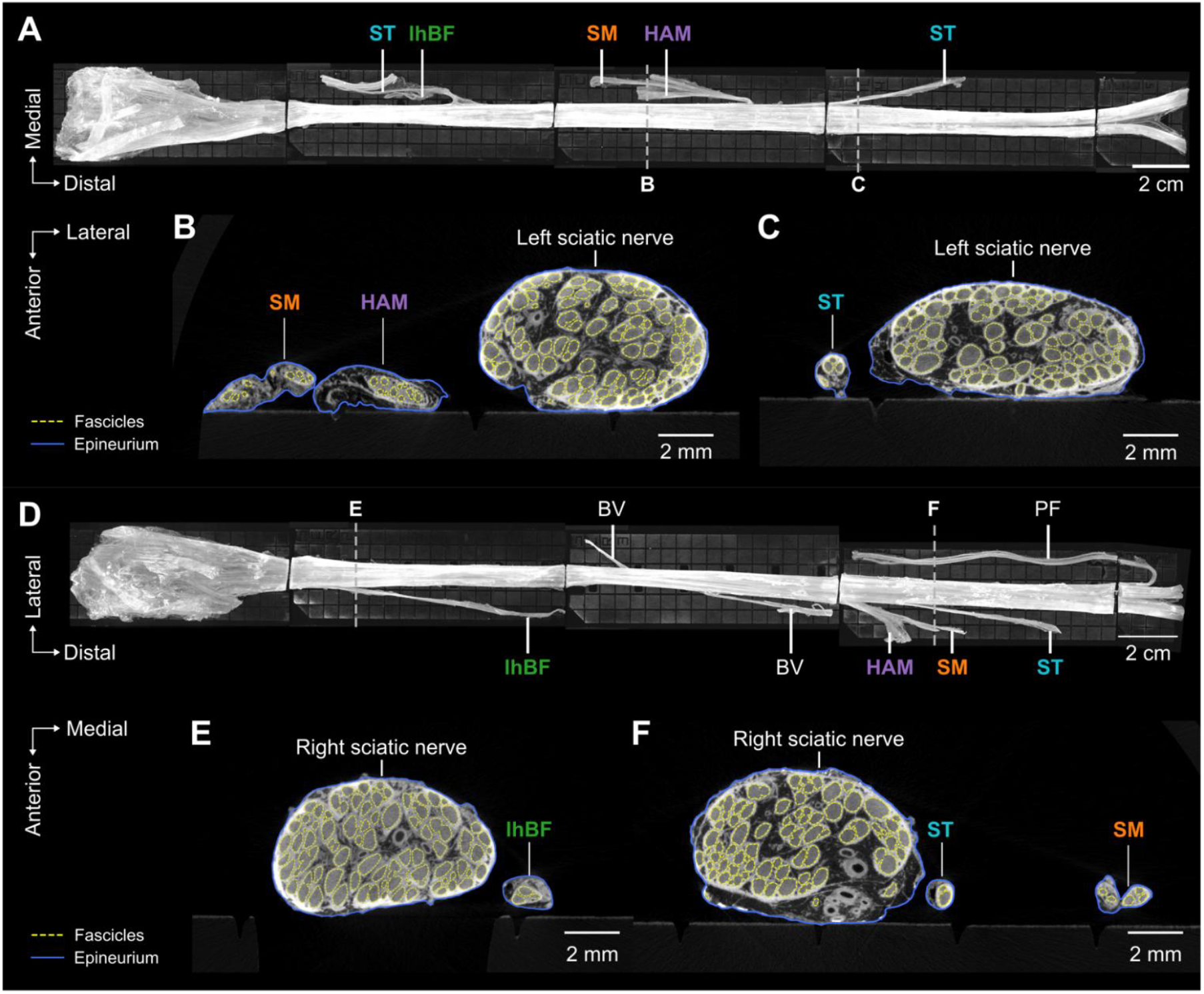
MicroCT imaging, segmentation, and identification of hamstring branches along bilateral sciatic nerves. The stitched maximum intensity projections of microCT images of left (A) and right (D) sciatic nerves are shown with branch targets annotated. (B, C, E, F) MicroCT cross sections along the left (B, C) and right (E, F) sciatic nerves at positions indicated by dashed lines in panels A and D showing fascicle (yellow dashed lines) and epineurium (blue lines) segmentations with labeled hamstring-innervating branches. BV, blood vessel; lhBF, long head of biceps femoris; HAM, hamstring part of the adductor magnus; PF, popliteal fossa; shBF, short head of biceps femoris; SM, semimembranosus; ST, semitendinosus.

Based on the segmentation, morphological measurements of the sciatic nerve and its fascicles were obtained along the ∼25 cm length of each side (Figure 5A, B). The nerve diameter was similar between the left and right sciatic nerves (mean: 6.46 mm, SD: 0.42 mm for left; mean: 6.78 mm, SD: 0.37 mm for right), and both sides reached minimum values at the level of the lesser trochanter. The left and right sciatic nerves had similar mean fascicle counts (mean: 82.52, SD: 9.35 for left; mean: 84.57, SD: 7.34 for right), and the number of fascicles on both sides peaked at ∼3 cm inferior to the lesser trochanter. Despite variations in fascicle count along the nerve, the mean and range of fascicle diameters remained consistent throughout the 25 cm length on both sides (mean: 0.40 mm, SD: 0.02 mm for left; mean: 0.41 mm, SD: 0.02 mm for right).

**Figure 5.**
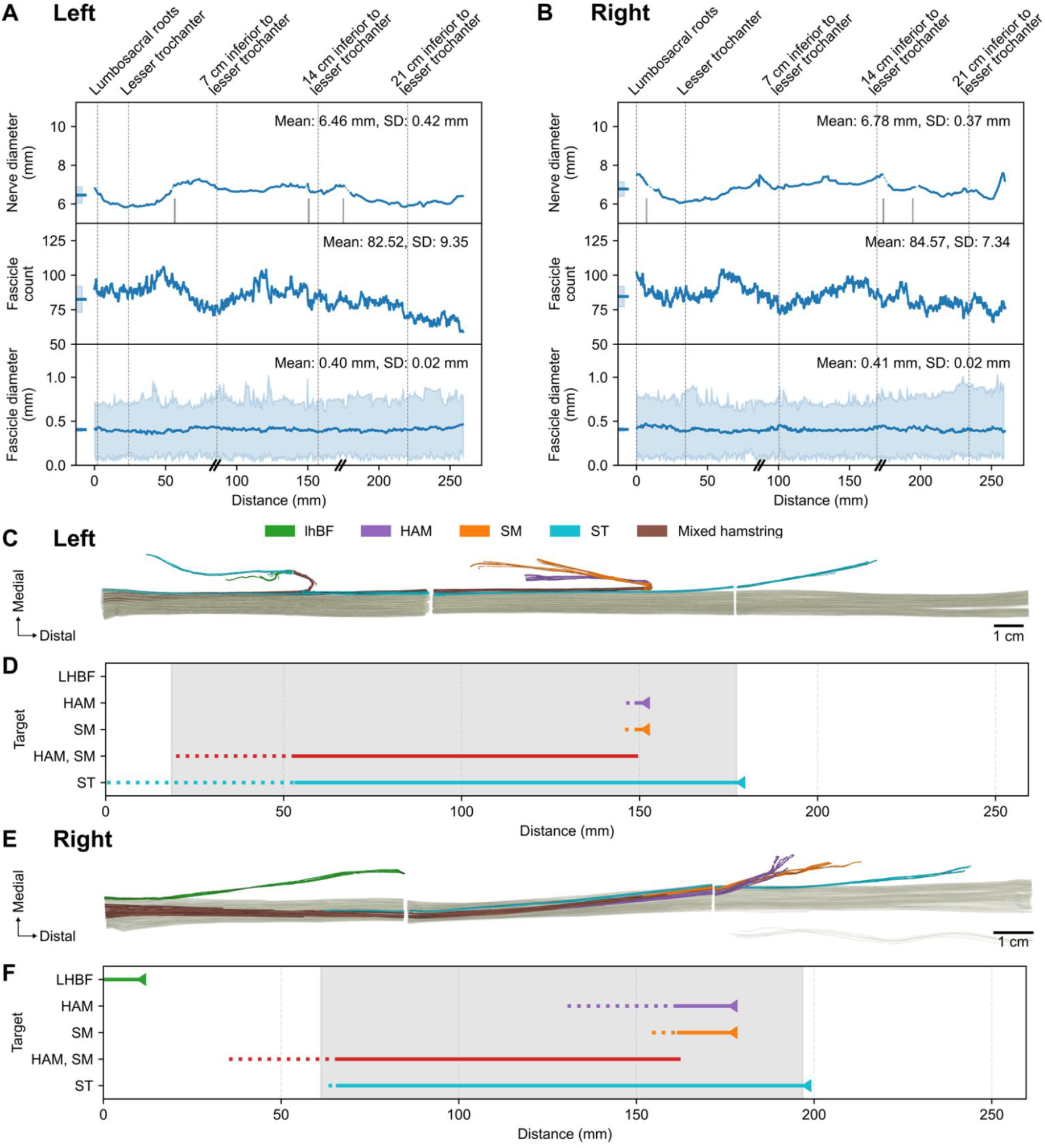
Quantified nerve morphology and fascicle tracing in the bilateral sciatic nerves. (A, B) Morphological measurements along the left (A) and right (B) sciatic nerves showing nerve diameter (top), fascicle count (middle), and fascicle diameter (bottom; line demarks median and shading demarks the range). Vertical dotted lines indicate the axial levels of anatomical landmarks. Mean and standard deviation values are provided for each metric (ticks with shading at x<0 and overlaid text in each top-right corner). In the top row (“Nerve diameter” plots), vertical gray ticks along the x axes indicate locations of branch points, and dashed blue lines indicate regions where the nerve diameter cannot be measured due to overlapping with branches. Double-slash marks (//) on the x-axes indicate transitions between microCT scanning grids. (C, E) 3D segmentation of fascicles along the left (C) and right (E) sciatic nerves, with hamstring fascicles color-coded by target muscles. Panel E is flipped relative to the raw image (Figure 4D) to show the anterior aspect of the nerve. (D, F) Hamstring fascicle tracing along the left (D) and right (F) sciatic nerves. Triangles indicate branch points. Solid line segments denote regions in which the traced fascicles from that branch remain distinct, without merging with fascicles from other targets (including other hamstring targets). Dotted line segments denote regions in which at least one fascicle remains distinct. Shaded regions denote length over which fascicles targeting one or more hamstring muscles remain distinct from fascicles targeting non-hamstring branches. lhBF, long head of biceps femoris; HAM, hamstring part of the adductor magnus; shBF, short head of biceps femoris; SM, semimembranosus; ST, semitendinosus.

### 3.3. Somatotopic Organization of Hamstring Fascicles Along and Within the Sciatic Nerve

Tracing from distal branch points proximally into the sciatic nerve, fascicles supplying hamstring muscles remained distinct over extended distances (Figure 5C–F). All ST fascicles remained distinct for 12.4 cm (left) and 13.1 cm (right); more proximally, at least one ST fascicle remained distinct over additional lengths of 5.3 cm (left) and 2.6 cm (right). HAM/SM fascicles shared the same distal branch points on both sides; all HAM/SM fascicles remained distinct for 9.6 cm (left) and 9.5 cm (right), after which at least one HAM/SM fascicle remained distinct over additional lengths of 0.32 cm (left) and 0.29 cm (right). lhBF fascicles remained distinct for 1.34 cm on the right side and merged with ST fascicles before reaching the sciatic on the left side. Fascicles targeting one or more hamstring muscles remained anatomically distinct from non-hamstring fascicles over continuous lengths of 15.9 cm (left) and 13.6 cm (right). The most proximal locations at which these hamstring fascicles remained distinct were 0.6 cm proximal to (left) and 2.7 cm distal to (right) the level of the lesser trochanter.

Hamstring fascicles showed a consistent cross-sectional organization within the sciatic nerves (Figure 6). Left hamstring fascicles tended to maintain a medial position throughout the sciatic nerve (Figure 5C), whereas right hamstring fascicles transitioned from an anterior position distally to a more medial position proximally (Figure 5E). On both sides, hamstring fascicles entered the sciatic nerve through medial branches and occupied medial or anteromedial positions within the nerve cross section (Figure 6A, B). As they progressed proximally toward the lumbosacral roots, these fascicles were predominantly localized to the anteromedial region. Morphological analysis of hamstring fascicles showed consistent mean diameters between sides (left: 0.38 mm, SD: 0.09 mm; right: 0.35 mm, SD: 0.06 mm), comparable to other sciatic nerve fascicles (Figure 6C, D). The left side contained fewer hamstring fascicles (mean: 6.91, SD: 3.80) than the right (mean: 9.16, SD: 4.17).

**Figure 6.**
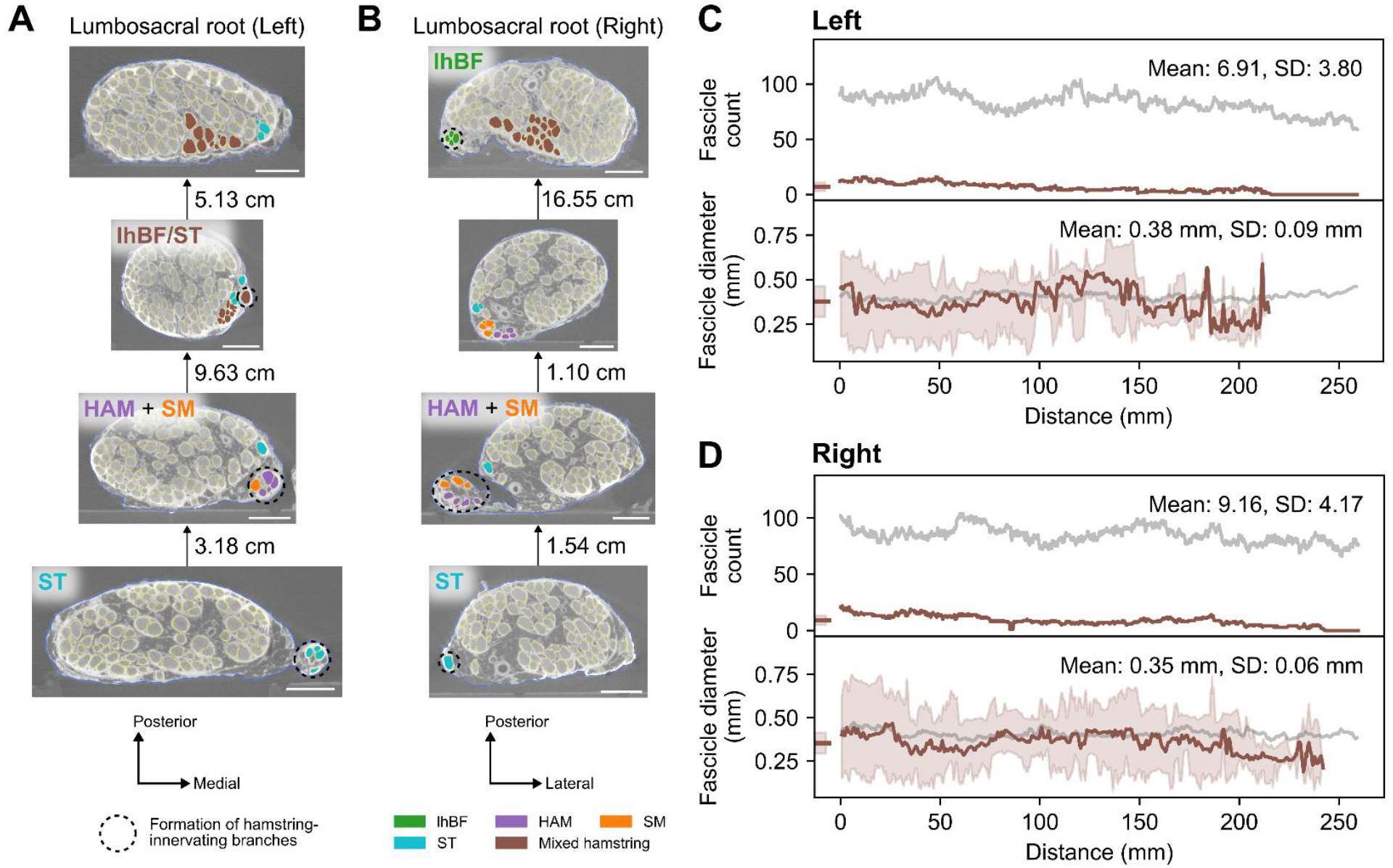
Somatotopic organization and quantified morphology of hamstring fascicles along bilateral sciatic nerves. (A, B) Fascicle maps along left (A) and right (B) sciatic nerves showing the tracked organization of hamstring-innervating fascicles from the most distal hamstring branch to the lumbosacral root level on each side. Fascicle masks are color-coded by their muscle targets; those targeting multiple hamstring muscles are categorized as mixed hamstring fascicles. Vertical arrows between microCT cross sections point from distal to proximal, with labeled inter-slice distances. Black dashed circles indicate the formation of branches targeting specific hamstring muscles labeled in the same cross section. In the second-to-top panel of A, lhBF and ST fascicles merged before entering the main trunk; the other ST fascicle originated from a more distal branch. Scale bars, 2 mm. (C, D) Morphological measurements comparing hamstring fascicles (brown) to all fascicles (gray) of the left (C) and right (D) sciatic nerves across ∼25 cm, showing fascicle count (top) and fascicle diameters (bottom; lines show the median fascicle diameters, and the shading shows the range). Mean and standard deviation values for the hamstring fascicles are provided for each metric (ticks with shading at x<0 and overlaid text in each top-right corner). lhBF, long head of biceps femoris; HAM, hamstring part of the adductor magnus; shBF, short head of biceps femoris; SM, semimembranosus; ST, semitendinosus.

## 4. Discussion

This work represents the first application of microCT imaging to comprehensively map the morphology and somatotopic organization of hamstring fascicles within human sciatic nerves, directly addressing a critical barrier to developing effective standing neuroprostheses. This novel approach visualizes the nerve’s internal structure in 3D at 11.4 μm isotropic resolution and enables tracking and measurement of fascicles across sciatic pathways over a range of 25 cm. In the proof-of-concept analysis of a single cadaver, hamstring fascicles were localized to the anteromedial region of the sciatic nerve and remained in functionally distinct groups for distances up to 13 cm. The microCT-derived data resolve anatomical features of complex neural pathways that were previously impossible to map at comparable resolution and range. The observed fascicular organization suggests that selective targeting of hamstrings may be feasible over proximal sciatic regions. This information, along with gross and fascicular morphology, provides anatomical input for modeling and optimizing neural interface design, stimulation parameters, and electrode placement.

### 4.1. Methodological Advantages of MicroCT-Based Fascicular Mapping

We performed detailed fascicular mapping and morphological measurements using microCT-derived data for bilateral sciatic nerves from a pilot subject. This approach presents a promising alternative to obtain data typically acquired through histological methods.

Traditional histological mapping, while considered the gold standard since the early 1900s (Raimondo et al., 2009), has significant limitations. Previous attempts to examine the proximal sciatic nerve using histological mapping (Gustafson et al., 2012) were discontinued due to the prohibitive cost and effort. The large intervals between slices (5–10 mm) prevent reliable tracking of actively mixing fascicles (i.e., the “plexiform” structure) in proximal nerve regions (McKinley, 1921; Stewart, 2003; Sunderland and Ray, 1948), forcing researchers to extrapolate fascicular grouping rather than directly observing changes in functional organization (Gustafson et al., 2012; McKinley, 1921; Sunderland and Ray, 1948). While computer-aided image reconstruction can accelerate the visualization of fascicle tracks within the nerve, it does not eliminate the need for labor-intensive sectioning and tissue processing (Zhong et al., 2015). Other potential imaging alternatives based on magnetic resonance (Yao et al., 2018) and ultrasound (Moayeri et al., 2010) cannot match the spatial resolution necessary for individual fascicle visualization. These methods, with resolutions of 0.2–0.5 mm, face challenges in resolving boundaries of individual fascicles (diameter ∼0.4 mm) to produce precise tracking data.

Overcoming these limitations, microCT imaging combines high resolution (∼10 μm) with extensive scanning range (∼10 cm) while preserving the longitudinal organization of neural tissue (Stauber and Müller, 2008). The high-resolution 3D visualization enables reliable, continuous tracking of detailed fascicular organization without sampling gaps (Thompson et al., 2020; Upadhye et al., 2022). It provides a transverse field of view of ∼3.5 cm, sufficient to capture entire nerve cross sections plus peripheral branches. When combined with segmentation and tracking algorithms (Buyukcelik et al., 2023; Sladjana et al., 2008), this integrated approach enables highly efficient fascicle-level mapping of complex neural pathways like the sciatic nerve that was previously unfeasible. Further, following microCT, tissues can undergo histology to provide fiber-level data (Upadhye et al., 2025a) for further structural and functional anatomical insights and to parameterize computational models.

### 4.2. Morphological Findings

We quantified and visualized the hamstring innervation pattern and fascicular morphology of the human sciatic nerve. Our findings generally align with results from prior studies while providing additional quantitative measurements along the nerve’s length.

#### 4.2.1. Branching Patterns and Nerve Morphology

The observed hamstring branches showed the expected medial branching points and trajectories (Figure 2), with each muscle receiving innervation from one or two branches, as previously reported (Bretonnier et al., 2019; Gustafson et al., 2012). The proximal-to-distal order of hamstring branches (lhBF, SM/HAM, ST) corresponds with the established pattern with only minor deviations (Gustafson et al., 2012). As previously reported (Gustafson et al., 2012), SM and HAM shared a common proximal branch trunk. Gross measurements confirmed the sciatic’s elliptical cross-sectional shape throughout its course (Figure 3). The average ratio between medial-lateral and anterior-posterior diameters (∼1.6) was smaller than in a prior study with ratios of major/minor axes (upper sciatic: 2.7, distal sciatic: 1.9) (Gustafson et al., 2012). Notably, the sciatic nerve narrowed to a more circular shape (aspect ratio ∼1) with minimum diameters (∼6 mm) bilaterally near the lesser trochanter of the femur (Figure 3 and Figure 5A, B).

During gross dissection, small vascular branches can be misidentified as nerve branches (Figure 2). MicroCT imaging identified two suspected cases of such misidentification in the present analysis (Figure 4).

#### 4.2.2. Fascicular Morphology

MicroCT analysis showed approximately constant fascicle diameters along the nerve length, averaging ∼0.4 mm, consistent with published values (∼0.42 mm) (Sladjana et al., 2008). The mean total fascicle count of 83 on both sides is consistent with an earlier observation (∼40–201) (Sunderland and Ray, 1948). Fascicle counts showed a distinct pattern along the nerve, reaching their maximum value (∼100) at 3–4 cm inferior to the lesser trochanter before decreasing to a local minimum at 7 cm inferior to the lesser trochanter (Figure 5A, B). The mean diameter of hamstring-innervating fascicles (0.36 mm) was slightly smaller than the overall sciatic nerve fascicles (∼0.4 mm) (Figure 6C, D). We observed modest laterality with fewer hamstring fascicles on the left side (∼7) compared to the right (∼9) (Figure 6C, D).

Tissue staining and extended microCT scanning sessions can cause tissue shrinkage and deformation due to dehydration and chemically induced degradation. While there is an inherent trade-off between enhanced contrast and tissue structural changes from staining (Descamps et al., 2014), these effects may be accounted for through calculated shrinkage factors (Buytaert et al., 2014) and improved humidity control in the scanning chamber.

### 4.3. Implications for Neuroprostheses Development

Realistic nerve and fascicular morphology obtained from microCT analysis provide the foundation for model-based design of implanted neuroprostheses to restore function. Nerve diameter directly informs electrode size which ensures a snug fit without inducing excessive tissue pressure (Freeberg et al., 2017; Naples et al., 1988) or blocking intrafascicular blood flow (Charkhkar et al., 2019). Fascicle diameters have important effects on the electric field and resulting neural responses: specifically, nerve fibers in smaller fascicles have lower activation thresholds (Davis et al., 2023).

The structural and functional organization of fascicles innervating target muscles determines the strategy needed for achieving selective stimulation (Charkhkar et al., 2019; Davis et al., 2023). For lower limb functions, the flat interface nerve electrode (FINE) (Tyler and Durand, 2002) was designed to target knee extensors by reshaping the femoral nerve to reduce fascicle-electrode distance, especially for centrally located fascicles. This led to improved ability to isolate activation of the individual quadriceps muscles with monopolar stimulation without field steering or other methods to achieve selectivity (Gustafson et al., 2009; Schiefer et al., 2010, 2008). However, our mapping suggests that hamstring fascicles clustered in the medial and anteromedial compartment of the sciatic, where the original FINE design may not provide sufficient contact density in an anatomically appropriate configuration to ensure selectivity and generalizability across the natural variations across potential recipients. The anatomical differences in fascicular organization highlight the need for modified or alternative configurations tailored to target hamstring fascicles based on their realistic arrangement within the nerve.

Our study suggests that fascicles innervating specific hamstring muscles can be tracked for considerable distances (e.g., up to 13 cm for ST). This finding aligns with early observations from serial histological cross sections (McKinley, 1921) and provides significantly more continuous and confident tracking. Notably, these distances fall within surgically accessible locations throughout the posterior thigh, particularly between the ischial tuberosity and the greater trochanter of the femur at the level of the gluteal fold. The distance over which fascicles remain distinct proximally to branch points offers valuable information for selecting more proximal intervention points while still achieving selective muscle activation. Given the tight anatomical constraints along the proximal sciatic nerve (Martin et al., 2015), this information complements the gross measurements of branch-free length and expands electrode placement options.

MicroCT imaging provides morphological data and fascicular organization along the entire sciatic nerve in 3D. This approach delivers more comprehensive anatomical information, capturing both intra- and inter-individual variations more effectively than previous histology-based methods. These detailed models thus enable optimization of electrode design and placement for optimal therapeutic effects. Our study demonstrates the microCT-based approach for mapping hamstring fascicles in the sciatic nerve using a single subject. While our findings provide valuable insights into targeting hamstring fascicles in standing neuroprostheses, they do not fully represent the anatomical variations across the population. Nevertheless, the improved throughput and accuracy of our fascicular mapping approach make it well-suited for application to larger cohort studies, which would enable systematic quantification of sciatic nerve anatomy and its variations. Furthermore, this methodology is not limited to the sciatic nerve and can be broadly applied to other peripheral nerves and functionally important fascicle groups.

## 5. Conclusions

We developed a novel approach to map the morphology and fascicular organization of hamstring fascicles in the human sciatic nerve using high-resolution microCT imaging. Our microCT-based methodology advances the ability to quantify the 3D morphology of complex neural pathways with greater efficiency and field of view than traditional histological methods while resolving fascicular morphology. The morphological data along the sciatic pathway—including nerve diameter, fascicle diameters, and fascicle count—and the spatial organization of hamstring-innervating fascicles provide critical anatomical information to inform selective neurostimulation. Future work will extend this approach to a larger cohort, mapping hamstring fascicles and characterizing inter-individual variability for clinical impact. We will also integrate these anatomical findings with realistic computational modeling to develop and validate novel neuroprostheses. This work establishes a methodological foundation for improving the selectivity and functional outcomes of motor and sensory lower limb neuroprostheses and for informing other orthopedic and neurosurgical interventions.

## 6. Acknowledgements

We thank the histology team of the Department of Biomedical Engineering of Case Western Reserve University (Jennifer Coleman, Aniya Hartzler) for their help with obtaining histological images. We thank Leina Lunasco, Anandakumar Shunmugavel, Jeya Shunmugavel, Elliot Crooks, Wonhee Han, Liam O’Reilly, and Victoria Zhao for their help performing preliminary cadaveric dissections, analyses, and background research. Funding was provided through the National Institute of Health (NIH) Stimulating Peripheral Activity to Relieve Conditions (SPARC) program (75N98022C00018), the Advanced Platform Technology (APT) Center, the US Department of Veterans Affairs (1I21RX005151), and the Department of Defense’s National Defense Science and Engineering Graduate (NDSEG) fellowship. The opinions expressed in this paper are those of the authors and do not reflect the views of the NIH, the Department of Health and Human Services, Department of Veterans Affairs, or the United States government.

## Supplementary Materials

### Supplementary Note 1

Data augmentation techniques applied in the network training.

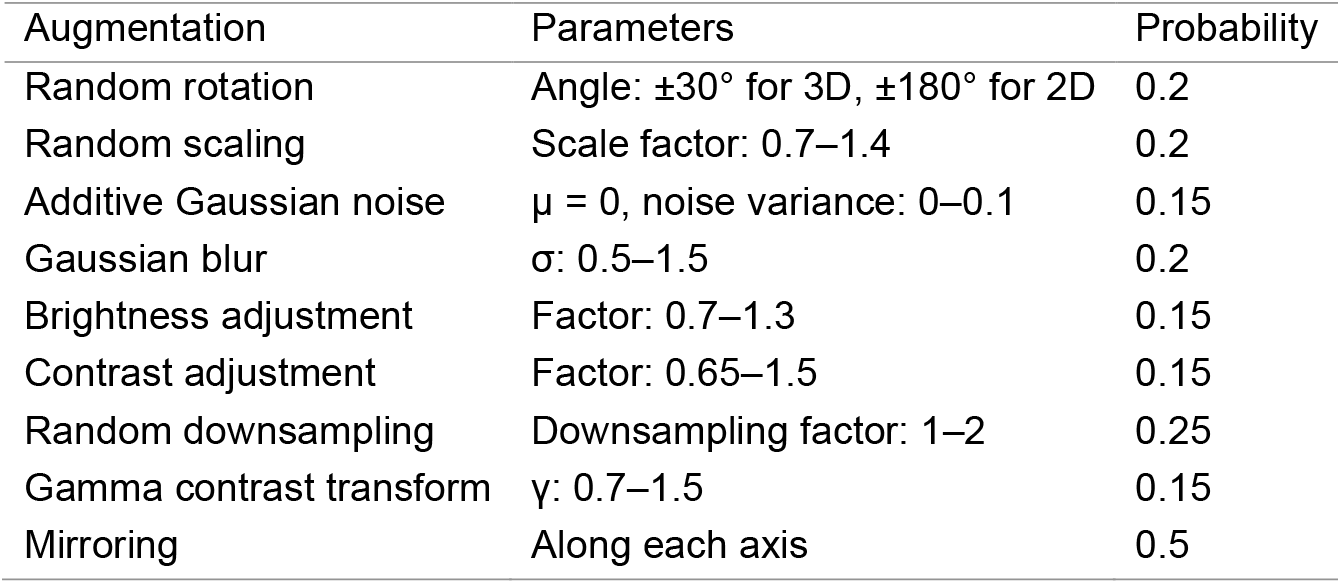

### Supplementary Note 2

In situ nerve diameter measurements.

**Table S1.** In situ nerve diameter measurements.

| Anatomical level | Side | Diameter (mm) |  | Aspect ratio |
| --- | --- | --- | --- | --- |
|  |  | Medial-lateral | Anterior-posterior |  |
| Formation of sciatic nerve from lumbosacral roots | Left | 7.2 | 6 | 1.2 |
|  | Right | 8.9 | 4.3 | 2.1 |
| Greater trochanter of femur | Left | 11.7 | 6.4 | 1.8 |
|  | Right | 11.7 | 3.7 | 3.2 |
| Lesser trochanter of femur | Left | 5.0 | 5.5 | 0.9 |
|  | Right | 5.3 | 4.3 | 1.2 |
| 7 cm inferior to lesser trochanter | Left | 6.8 | 4.4 | 1.6 |
|  | Right | 7.2 | 4.3 | 1.7 |
| 14 cm inferior to lesser trochanter | Left | 7.1 | 4.2 | 1.7 |
|  | Right | 6.5 | 5.1 | 1.3 |
| 21 cm inferior to lesser trochanter | Left | 6.8 | 3.9 | 1.7 |
|  | Right | 7.5 | 5.2 | 1.4 |

### Supplementary Note 3

Histological validation of fascicular structures identified in microCT images of representative regions of sciatic nerve and branch. We adopted a protocol used for histology of cadaveric human vagus nerves (Coleman et al., 2025). After microCT imaging, nerves were removed from the acrylic boards and sectioned into 15 mm segments. Samples were fixed and processed using a HistoCore PEGASUS Tissue Processor (Leica Biosystems, Deer Park, IL) at 45°C . Processing included an initial 30-min wash in 10% neutral buffered formalin, followed by graded ethanol dehydration with 70% ethanol (15 min) and progressing through increasing concentrations (15, 15, 1, and 4 minutes) to 100% ethanol, after which tissues underwent pressurized paraffin infiltration. The 15 mm-long segments were cut into 5 mm-long segments and embedded in paraffin. The tissue was sectioned into 4 µm slices using a HistoCore AUTOCUT microtome (Leica Biosystems, Deer Park, IL) and affixed to slides via a warm (36°C) distilled water bath. Slides were stained using the StatLab Select progressive H&E protocol (StatLab, McKinney, TX), which included hematoxylin and eosin application followed by graded alcohol dehydration and xylene clearing prior to coverslipping. Digital images of the stained slides were captured using a Zeiss Axioscan Z7 automatic slide scanner at 20x magnification.

**Figure S1.**
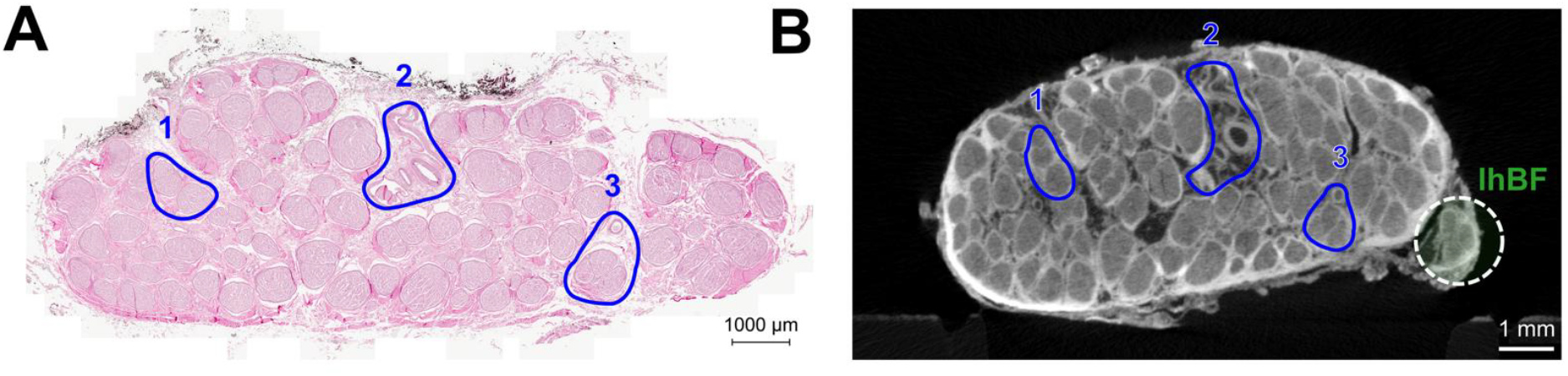
Example histological validation of microCT images of the right sciatic nerve at the lumbosacral roots. (A) Histological cross section stained with hematoxylin and eosin. (B) Corresponding microCT cross section at the same anatomical level. In both panels, blue outlines and labels indicate matched anatomical features (e.g., fascicle groups and blood vessels) identified across the two modalities. The long head of the biceps femoris branch (lhBF), highlighted with a dotted circle, is visible only in the microCT image.

**Figure S2.**
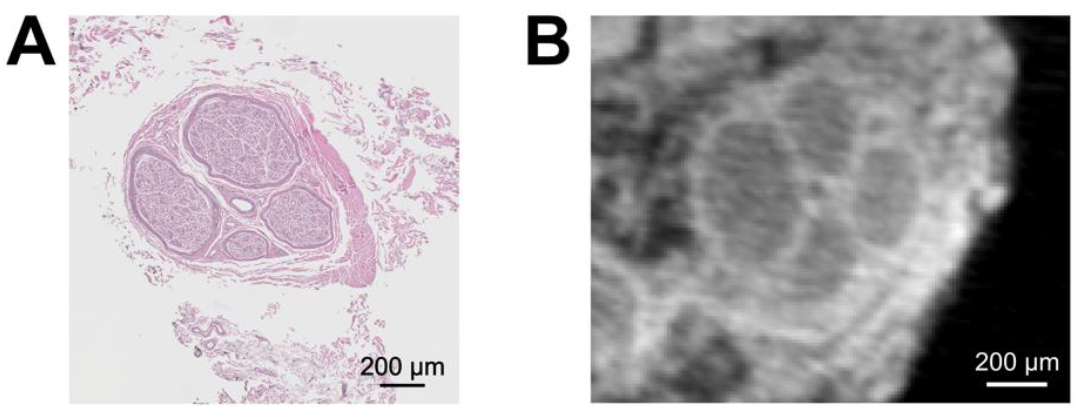
Example histological validation of microCT images of the right branch to the long head of biceps femoris (lhBF) muscle. (A) Histological slide with H&E stain showing the cross-sectional morphology of the right branch to lhBF. (B) MicroCT cross section showing the corresponding fascicular morphology of the same branch.

